# Complete chloroplast genomes of *Anthurium huixtlense* and *Pothos scandens* (Pothoideae, Araceae): unique inverted repeat expansion and contraction affect rate of evolution

**DOI:** 10.1101/2020.03.11.987859

**Authors:** Abdullah, Claudia L. Henriquez, Furrukh Mehmood, Monica M. Carlsen, Madiha Islam, Mohammad Tahir Waheed, Peter Poczai, Thomas B. Croat, Ibrar Ahmed

## Abstract

The subfamily Pothoideae belongs to the ecologically important plant family Araceae. Here, we report the chloroplast genomes of two species of the subfamily Pothoideae: *Anthurium huixtlense* (163,116 bp) and *Pothos scandens* (164,719 bp). The chloroplast genome of *P. scandens* showed unique inverted repeats (IRs) contraction and expansion, which increases the size of the large single copy (102,956) region and decreases the size of the small single-copy (6779 bp) region. This led to duplication of many single-copy genes due to transfer to IR regions from the small single-copy (SSC) region, whereas some duplicate genes became single copy due to transfer to large single-copy regions. The rate of evolution of protein-coding genes was affected by the contraction and expansion of IRs; we found higher mutation rates for genes that exist in single-copy regions as opposed to IRs. We found a 2.3-fold increase of oligonucleotide repeats in *P. scandens* when compared with *A. huixtlense*, whereas amino acid frequency and codon usage revealed similarities. We recorded higher transition substitutions than transversion substitutions. The transition/transversion ratio was 2.26 in *P. scandens* and 2.12 in *A. huixtlense*. We also found a higher rate of transversion substitutions linked with non-synonymous substitutions than synonymous substitutions. The phylogenetic inference of the limited species showed the monophyly of the Araceae subfamilies. Our study provides insight into the molecular evolution of chloroplast genomes in the subfamily Pothoideae and family Araceae.

## 1. Introduction

The plant family Araceae belongs to the order Alismatales. Araceae is a large and ancient monocot family and consists of 114 genera and 3750 species (Christenhusz and Byng 2016), although Boyce and Croat (2018) have estimated approximately 6500 species. It is the most diverse monocotyledon family in terms of morphology (Gunawardena and Dengler 2006) and ecological distribution (Cabrera et al. 2008). Species of Araceae have been subdivided into eight subfamilies and exist in tropical and temperate regions (Cabrera et al. 2008; Cusimano et al. 2011; Nauheimer et al. 2012). Pothoideae is the second largest subfamily, with approximately 1010 described species and approximately 2072 estimated species (Boyce and Croat 2018) in four genera, including *Pothos* L., *Pothoidium* Schott*, Pedicellarum* M.Hotta, and *Anthurium* Schott (Cabrera et al. 2008). The largest genera are *Pothos* and *Anthurium*, with approximately 57 and 950 described species and approximately 70 and 2000 estimated species, respectively (Boyce and Croat 2018). The other two genera, *Pothoidium* and *Pedicellarum*, are monospecific.

The chloroplast is a self-replicating organelle that plays a vital role in photosynthesis and in the synthesis of fatty acids and amino acids (Cooper 2000). In most plant lineages, the chloroplast contains its own circular double-stranded genome and has a primarily quadripartite structure in which a pair of long inverted repeat regions (IRa and IRb) separate the large single-copy (LSC) and small single-copy (SSC) regions (Palmer 1985). However, linear chloroplast genomes have also been reported (Oldenburg and Bendich 2016) in some species. Moreover, a quadripartite structure has not been observed in the chloroplast genomes of various species, such as Pinaceae (Wu et al. 2011), Cephalotaxaceae (Yi et al. 2013), and Taxodiaceae (Hirao et al. 2008). The size of the chloroplast genome of photosynthetic plants varies from 107 kb (*Cathaya argyrophylla* Chun & Kuang) to 218 kb (*Pelargonium × hortorum* L.H.Bailey) (Daniell et al. 2016). Chloroplast genomes are inherited from a single parent and show significant polymorphism (Daniell 2007; Daniell et al. 2016), which makes them well-suited for studies on phylogenetics and population genetics (Ahmed et al. 2013; Ahmed 2014; Henriquez et al. 2014).

Despite a relatively conserved structure, including gene organization, gene content, and intron content within genes (Iram et al. 2019; Mehmood et al. 2020; Shahzadi et al. 2020), chloroplast genomes have also undergone gene loss, intron loss, gene rearrangement, pseudogenization, gene duplication, and uneven expansion and contraction of IR regions. These events have led to a variable number of genes in the chloroplast genomes of angiosperms (Menezes et al. 2018; Abdullah et al. 2020; Henriquez et al. 2020a). Moreover, the shifting of genes to single-copy regions from IR or vice versa due to IR contraction and expansion also affect the rate of DNA sequence evolution. The phenomenon is known as rate heterotachy (Lockhart et al. 2006). Previous studies of subfamilies Lemnoideae and Aroideae revealed unique and uneven contraction and expansion of IR regions, which led to a variable number of genes and gene rearrangements in the chloroplast genomes of several of their respective taxa (Wang and Messing 2011; Choi et al. 2017; Kim et al. 2019; Henriquez et al. 2020a). The aforementioned studies did not include species of the subfamily Pothoideae.

In this study, a comparison of the *de novo* assembled chloroplast genomes of *A. huixtlense* Matuda and *P. scandens* L. with chloroplast genomes of other Araceae species confirmed unique events of IR contraction and expansion in the chloroplast genome of *P. scandens*. The results reveal the transfer of IR genes to the LSC region at the junction of JLB (LSC/IRb) and the transfer of SSC genes (except *rps15* and *ycf1*) to the IR region at the junction of JSB (IRb/SSC). This transfer promotes heterotachy in Pothoideae by affecting the rate of evolution of these genes. To the best of our knowledge, this shortening of the SSC region due to unique IR contraction and expansion and the effects on gene evolution rate are reported here in Araceae for the first time. These results improve our understanding of the evolution of chloroplast genomes in Araceae.

## 2. Materials and Methods

### 2.1 DNA extraction and sequencing

We collected fresh healthy leaves of *P. scandens* and *A. huixtlense* from the Aroid Greenhouse at the Missouri Botanical Garden in St. Louis, Missouri. Total genomic DNA was extracted from these leaves using a Qiagen DNeasy Minikit (Qiagen, Germantown, Maryland, USA) following Henriquez et al. (2020a). Confirmation of the quality and quantity of DNA was performed using 1% gel electrophoresis and Nanodrop (ThermoScientific, Delaware, USA). Library preparation and sequencing were performed using TruSeq kits (Illumina, Inc., San Diego, California) in the Pires lab at the University of Missouri, Columbia following Henriquez et al. (2020a).

### 2.2 *De novo* assembly and annotation

The quality of raw reads was analyzed by FastQC (Andrews 2017) and MultiQC (Ewels et al. 2016) for comparison. After quality confirmation, the Fast-Plast v. 1.2.2 pipeline (https://github.com/mrmckain/Fast-Plast) was initially used to assemble the raw reads following similar parameters previously employed for the assembly of chloroplast genomes of subfamilies Aroideae and Monsteroideae (Henriquez et al. 2020b, a). The resulting assembly from Fast-Plast was further confirmed by *de novo* assembly using Velvet v.1.2.10 following Abdullah et al. (2019a, 2020) using Kmer values of 61, 71, and 81. Validation and coverage depth analyses of *de novo* assembled genomes were performed by mapping short reads to their respective assembled chloroplast genomes. The assembled chloroplast genomes were annotated using GeSeq (Tillich et al. 2017) and the circular diagrams of the annotated genomes were drawn using OrganellarGenomeDRAW (OGDRAW v.13.1) (Greiner et al. 2019). The five-column tab-delimited tables were generated for *de novo* assembled chloroplast genomes using GB2sequin (Lehwark and Greiner 2019) and were submitted to the National Center for Biotechnology Information (NCBI) under accession number MN046891 (*P. scandens*) and MN996266 (*A. huixtlense*). The raw reads were also submitted to the sequence read archive (SRA) of NCBI under BioProject number PRJNA547619.

### 2.3 Characterization and comparative analyses of chloroplast genomes

Characterization of the chloroplast genomes of *P. scandens* and *A. huixtlense* and analyses of amino acid frequency and codon usage were performed in Geneious R8.1 (Kearse et al. 2012). Oligonucleotide repeats were determined using REPuter (Kurtz et al. 2001) by setting the parameter of minimum repeat size ≥ 30 and with minimum repeat similarity of 90%.

The chloroplast genome structure and gene content of *P. scandens* and *A. huixtlense* were compared with eight previously reported chloroplast genomes, including *Anchomanes hookeri* (Kunth) Schott, *Anubias heterophylla* Engler, *Zantedeschia aethiopica* (L.) Spreng. (Henriquez et al. 2020a), *Epipremnum aureum* (Linden & André) G.S. Bunting (Tian et al. 2018), *Spathiphyllum kochii* Engl. & K. Krause (Han et al. 2016), *Spirodela polyrrhiza* (L.) Schleid., *Wolffiella lingulata* Hegelm. (Wang and Messing 2011), and *Symplocarpus renifolius* Schott ex Tzvelev (Choi et al. 2017). The gene content and rearrangement of the genome were compared by integrated Mauve alignment (Darling et al. 2004) in Geneious R8.1 based on collinear blocks analyses. IR contraction and expansion were studied among these species using IRscope (Amiryousefi et al. 2018a).

We also analyzed synonymous (K_s_) and non-synonymous (K_a_) substitutions and their ratio (K_a_/K_s_). *Symplocarpus renifolius*, a species from the early diverging subfamily Orontioideae, was used as a reference and 75 protein-coding genes of *P. scandens* and *A. huixtlense* were aligned to the protein-coding genes of *Symplocarpus renifolius* by MAFFT alignment (Katoh et al. 2005). These alignments were analyzed for the determination of K_s_ and K_a_ substitutions and Ka/Ks using DnaSP (Rozas et al. 2017) as in previous studies (Abdullah et al. 2019a, 2020; Henriquez et al. 2020b). We also determined the extent of transition and transversion substitutions that linked with Ks and Ka substitutions. For this purpose, we selected 11 genes from the genome of *P. scandens* that had various Ka/Ks values and analyzed the extent of transition and transversion types of substitutions with Ks and Ka substitutions in Geneious R8.1 (Kearse et al. 2012) following Abdullah et al. (2019a).

We also used *Symplocarpus renifolius* as a reference to analyze the effect of rate heterotachy on the evolution of protein-coding genes. We considered genes of LSC, SSC, and IR of *Symplocarpus renifolius* and determined the rate of evolution of the respective genes in the chloroplast genomes of *P. scandens* and *A. huixtlense*. We also separately determined the rate of evolution of protein-coding genes that transferred from IRs to LSC or from SSC to IR to elucidate the changes in evolution rate. We concatenated genes of each region and aligned using MAFFT (Katoh et al. 2005). The types of transition and transversion substitutions in *P. scandens* and *A. huixtlense* were also determined from the alignment of genes from LSC, SSC, and IR.

### 2.4 Phylogenetic inference

The phylogenetic tree was inferred using 26 species of Araceae, with *Acorus americanus* (Acoraceae) as the outgroup. The accession numbers of all included species are provided in Table S1. The complete chloroplast genomes, excluding IRa, were aligned by MAFFT (Katoh et al. 2005) and the phylogeny was inferred using the IQ-tree program (Nguyen et al. 2015; Kalyaanamoorthy et al. 2017; Hoang et al. 2018) with default parameters using a previous approach (Abdullah et al. 2019a, 2020).

## 3. Results

### 3.1 Assembly and characterization of chloroplast genomes

The sequencing of 100-bp single-end reads generated 3.69 GB data (14.13 million reads) for *A. huixtlense* and 5.8 GB data (22.2 million reads) for *P. scandens*. Whole-genome shotgun reads contained 0.22 million reads in *A. huixtlense* and 0.77 million reads of chloroplast origin in *P. scandens*. These chloroplast reads were used for *de novo* assembly and provided average coverage depths of 468× for *P. scandens* and 138× for *A. huixtlense*.

The sizes of the complete chloroplast genomes were 163,116 bp for *A. huixtlense* and 164,719 bp for *P. scandens*. The sizes of the LSC and SSC regions showed a high level of variation between the two species due to unique IR contraction and expansion in the *P. scandens* chloroplast genome (Table 1). The GC content was highest in IR regions, followed by LSC and SSC regions. A high level of variation exists in the GC content of the chloroplast genome of both species.

**Table 1.** Comparative analyses of chloroplast genomes of *P. scandens* and *A. huixtlense*

| **Characteristics** |  | ***Pothos scandens*** | ***Anthurium huixtlense*** |
| --- | --- | --- | --- |
| Size (base pair; bp) |  | 164,719 | 163,116 |
| LSC length (bp) |  | 102,956 | 89,260 |
| SSC length (bp) |  | 6779 | 22,982 |
| IR length (bp) |  | 27,492 | 25,437 |
| Number of genes |  | 135 | 130 |
| Protein-coding genes |  | 90 | 85 |
| tRNA genes |  | 36 | 37 |
| rRNA genes |  | 8 | 8 |
| Duplicate genes |  | 22 | 17 |
| GC content | Total (%) | 35.4 | 36.2 |
|  | LSC (%) | 34.7 | 34.5 |
|  | SSC (%) | 28.8 | 28.5 |
|  | IR (%) | 37.3 | 42.7 |
|  | CDS (%) | 37.4 | 38 |
|  | rRNA (%) | 54.9 | 55 |
|  | tRNA (%) | 53.3 | 53.2% |

We found 114 unique functional genes in both species, including 80 protein-coding genes, 30 tRNA genes, and 4 rRNA genes (Fig. 1A and Fig. 1B). The *inf*A gene was observed as a pseudogene in both species, whereas the *rpl*23 gene was observed as pseudogene in *P. scandens* due to the generation of an internal stop codon. The total number of genes varied between the two species due to IR contraction and expansion. We found 130 genes in *A. huixtlense*, including 37 tRNA genes, 85 protein-coding genes, and 8 rRNA genes. We also observed 17 genes that were duplicated in the IR regions in *A. huixtlense*, including 7 tRNA genes (2 genes also contain introns), 4 rRNA genes, and 6 protein-coding genes (3 genes also contain introns) (Fig. 1A). In *P. scandens*, we found 135 genes due to expansion of the IR region, including 36 tRNA genes, 90 protein-coding genes, and 8 rRNA genes (Fig. 1B). We found 22 genes that were duplicated in the IR regions in *P. scandens*, including 6 tRNA genes (2 genes also contain introns), 4 rRNA genes, and 12 protein-coding genes (2 genes also contain introns) (Fig. 1B).

**Fig. 1.**
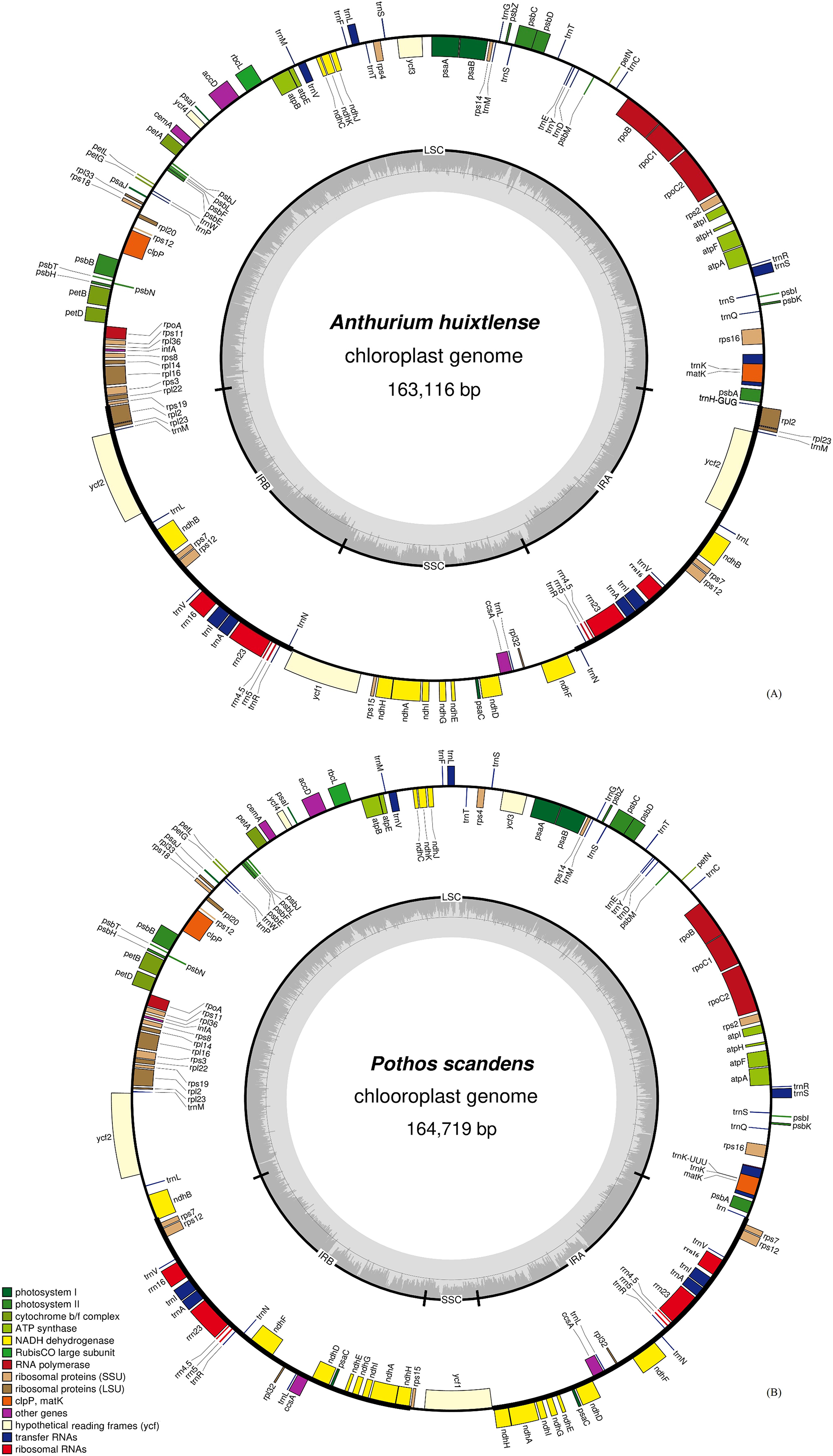
Circular diagram of chloroplast genomes of *A. huixtlense* and *P. scandens*. LSC, SSC, and IR represent large single copy, small single copy, and inverted repeat regions, respectively. The genes located inside are transcribed counterclockwise, whereas the genes located outside are transcribed clockwise.

### 3.2 Amino acid frequency and codon usage

The highest frequency observed was for leucine followed by iso-leucine, whereas the lowest frequency observed was for cysteine (Fig. 2). Relative synonymous codon usage (RSCU) analyses revealed high encoding frequency for codons containing A or T at the 3’ end and having an RSCU value of ≥ 1, whereas low encoding frequency was observed for codons having C or G at the 3’ and having RSCU < 1 (Table S2).

**Fig 2.**
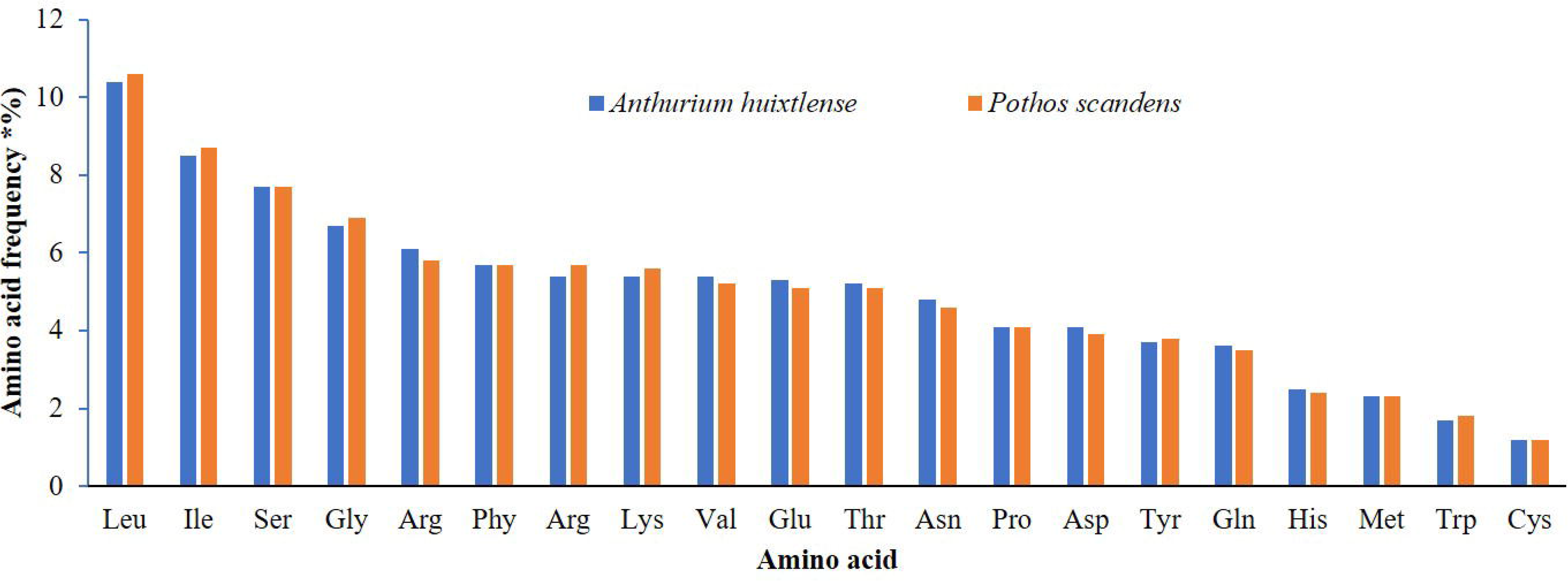
Amino acid frequency in *A. huixtlense* and *P. scandens*. The x-axis shows amino acids whereas the Y-axis shows percentage of amino acid frequency.

### 3.3 Repeats analyses

REPuter detected four types of oligonucleotide repeats in the chloroplast genomes of *A. huixtlense* and *P. scandens*. The number of repeats and types varied in both species to a high degree. We observed 37 repeats in *A. huixtlense* and 85 repeats in *P. scandens*. We observed 9 forward, 12 palindromic, 6 complementary, and 10 reverse repeats in *A. huixtlense*. In *P. scandens* we observed 21 forward, 33 palindromic, 8 complementary, and 23 reverse repeats (Fig. 3A). Most of the repeats were found in LSC regions instead of SSC and IR regions (Fig. 3B). Most of the repeats ranged in size from 40 bp to 44 bp in *A. huixtlense*. In *P. scandens*, most of the repeats varied in size from 35 bp to 39 bp (Fig. 3C). Details are provided in Table S3.

**Fig. 3.**
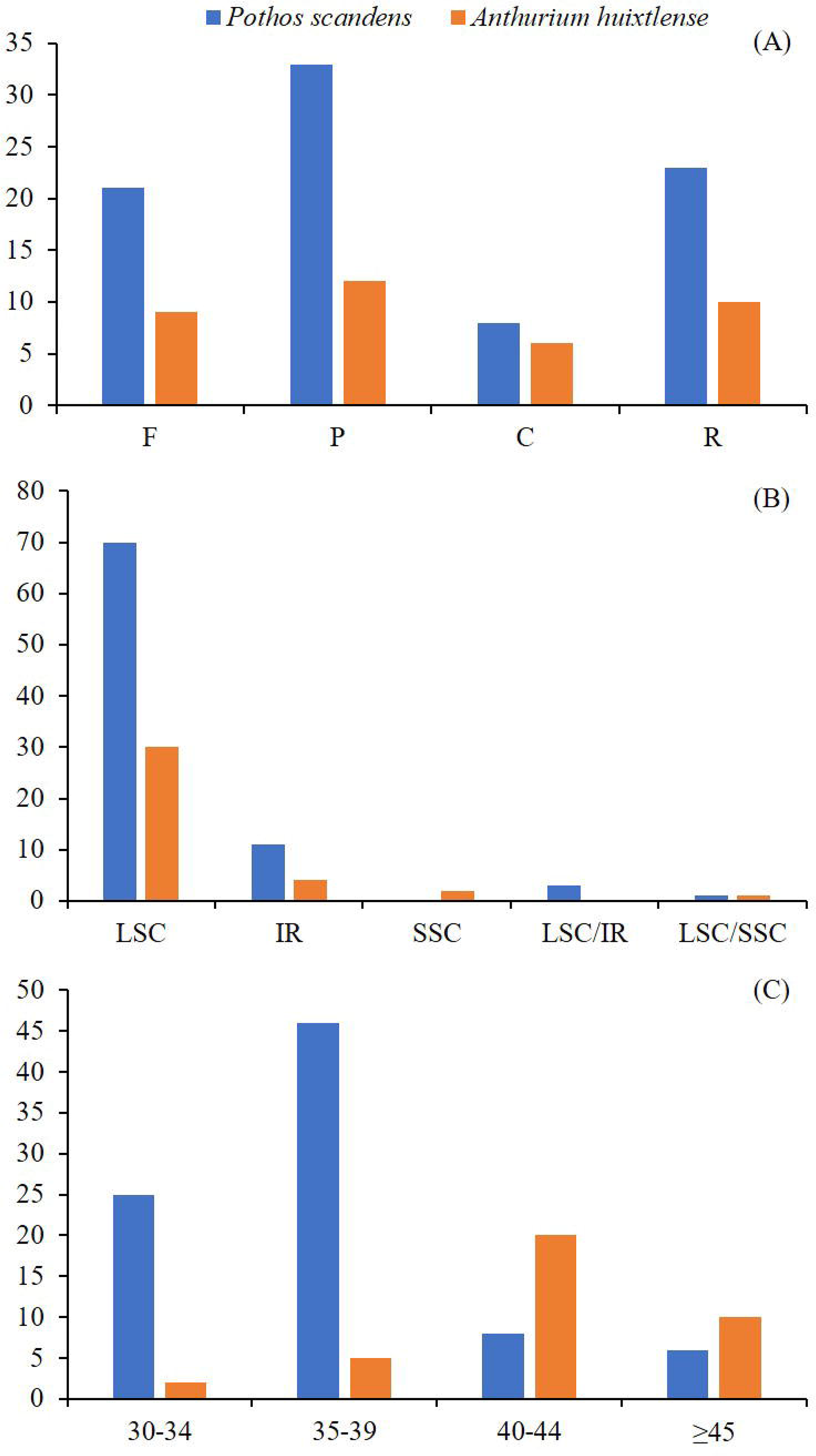
Analyses of repeats in *A. huixtlense* and *P. scandens*. (A) represents types of repeats. (B) Distribution of repeats in the chloroplast genomes (C). The size of repeats in the genome. F = Forward, C = complementary, R = reverse, and P = palindromic, LSC = large single copy, SSC = small single copy, IR = inverted repeat, LSC/SSC and LSC/IR represent those repeats shared between regions.

#### Transition and transversion substitutions and evolution rate of protein-coding genes

The evolution rate of protein-coding genes revealed strong purifying selection on these genes and that none of the genes are under positive selection pressure. Except for a few genes that showed neutral selection, all other genes showed purifying selection (Table S4) (average Ks = 0.16, Ka = 0.026, and Ka/Ks = 0.18). As expected, the highest purifying selection pressure was observed for genes that are involved in photosynthesis.

In the protein-coding genes of *P. scandens*, we found 4061 substitutions. Of these, 2814 contained transition substitutions (Ts) and 1247 contained transversion substitutions (Tv); the Ts/Tv ratio was 2.26. In *A. huixtlense*, we recorded 3960 substitutions, of which 2690 were Ts and 1270 were Tv (Ts/Tv = 2.12) (Table 2). These values revealed higher Ts than Tv. Examination of 11 protein-coding genes revealed a Ts/Tv of 2.79 for synonymous substitutions and a Ts/Tv of 1.43 for non-synonymous substitutions. Hence, a high number of Tv leads to non-synonymous substitutions as compared to Ts, which leads to synonymous substitutions.

**Table 2.**
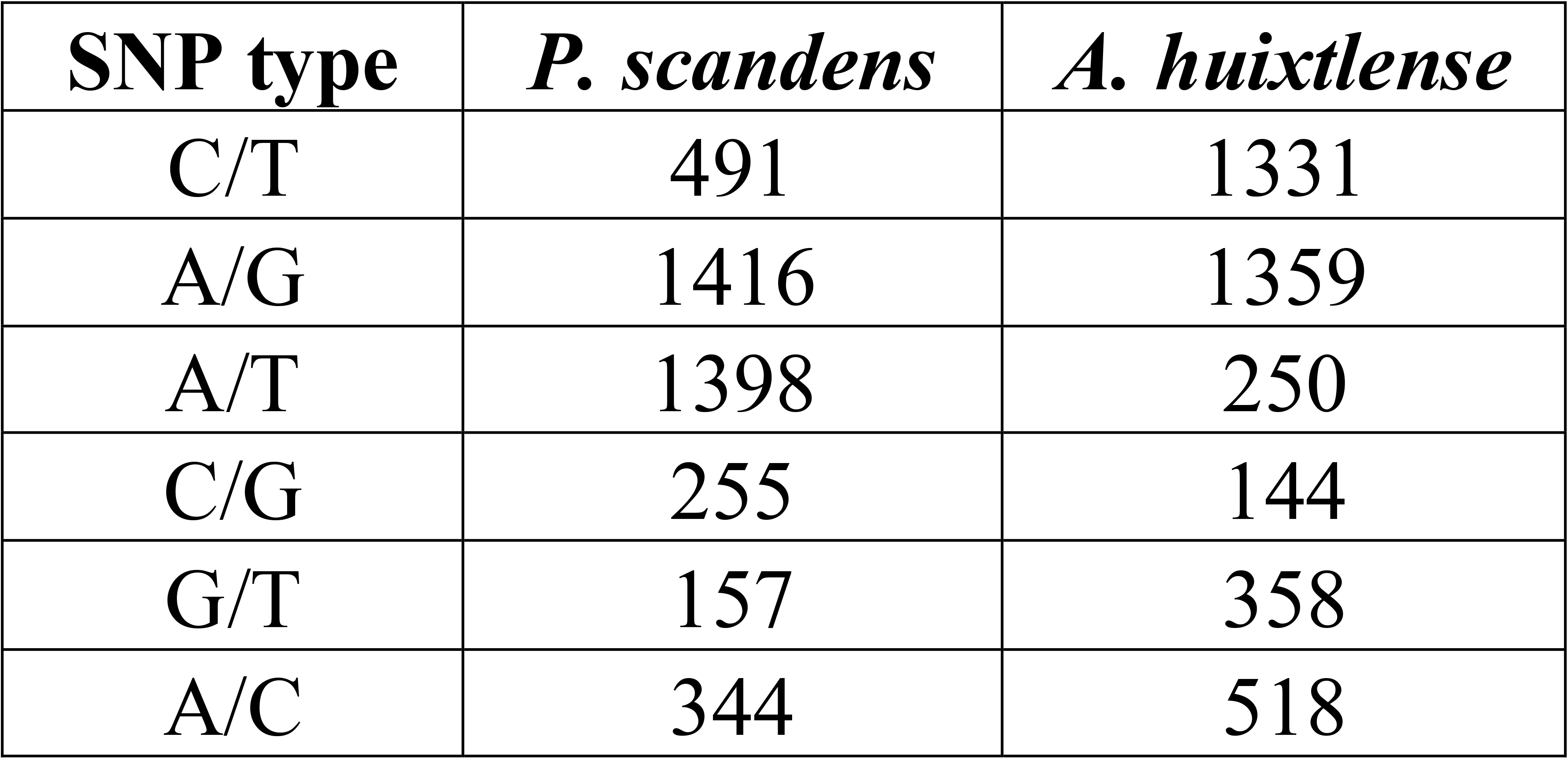
Transition and transversion substitutions in coding genes of *P. scandens and A. huixtlense*

### 3.4 Gene rearrangement and inverted repeats contraction and expansion

The genomes of Pothoideae show unique gene and structural rearrangements. The *P. scandens* chloroplast genome showed unique IR contraction and expansion, which led to a variable number of genes and also a change in gene arrangement. At the junction of JLB (LSC/IRb), the contraction of IR leads to expansion of the LSC region, whereas at the JSB (IRb/SSC) junction, the expansion of IR leads to contraction of the SSC region. Hence, many genes (*rpl*2, *rpl*23, *trn*M, *ycf*2, *trn*L, *ndh*B) that are usually duplicated in the IRs become single copy at the junction of JLB due to their transfer to LSC. In contrast, many genes (*ndh*H, *ndh*A, *ndh*I, *ndh*G, *ndh*E, *psa*C, *ndh*D, *ccs*A, *trn*L, *rpl*32, and *ndh*F) that usually exist as single copy due to their existence in SSC exist in duplicate due to their transfer to IR regions (Fig. 1). The arrangement of genes in LSC in both *A. huixtlense* and *P. scandens* did not change due to contraction of IR regions and gene arrangement was found to be similar to other species (*Spathiphyllum kochii*, *Epipremnum aureum*, *Symplocarpus renifolius*, and *Anubias heterophylla*), as shown in Colinear block of Mauve alignment (Fig. 4). However, the genes of the SSC region showed variation in gene arrangement (Fig. 4). In the genome of *A. huixtlense*, the SSC was inverted when compared to other species of Aroideae. However, this could not be considered any important evolutionary events as chloroplast genome exist in two equimolar states and can be differed by orientation of SSC region (Walker et al. 2015).

**Fig. 4.**
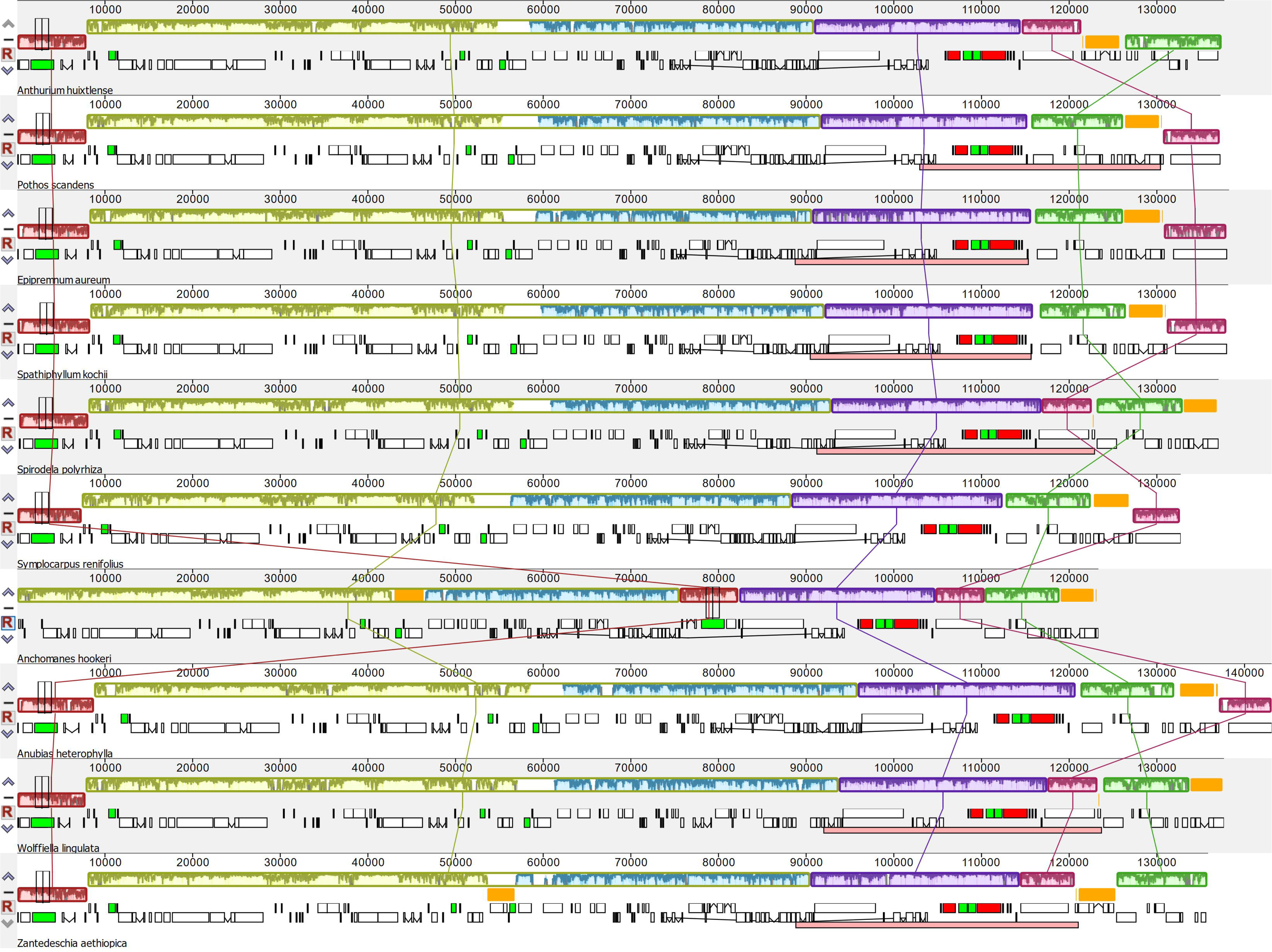
Analyses of gene arrangements among the species of Araceae based on Mauve alignment. White boxes indicate protein-coding genes, red indicate rRNA, black indicate tRNA, green indicate intron-containing tRNA, the line between two white boxes indicates intron-containing genes.

The contraction and expansion of IR regions at the junctions JLB (LSC/IRb), JSB (IRb/SSC), JSA (SSC/Ira), and JLA (IRa/LSC) were analyzed among the species of Araceae. We observed five types of variation in the junctions (Fig. 5). Type A included *P. scandens*, type B included *A. huixtlense*, *Epipremnum aureum* (see below), *Spathiphyllum kochii*, *Symplocarpus renifolius*, and *Anubias heterophylla*, type C included *Wolffiella lingulata* and *Spirodela polyrhiza*, type D included *Zantedeschia aethiopica*, and Type E included *Anchomanes hookeri*. These results show that the *P. scandens* chloroplast genome displays a novel type of IR contraction and expansion.

**Fig. 5.**
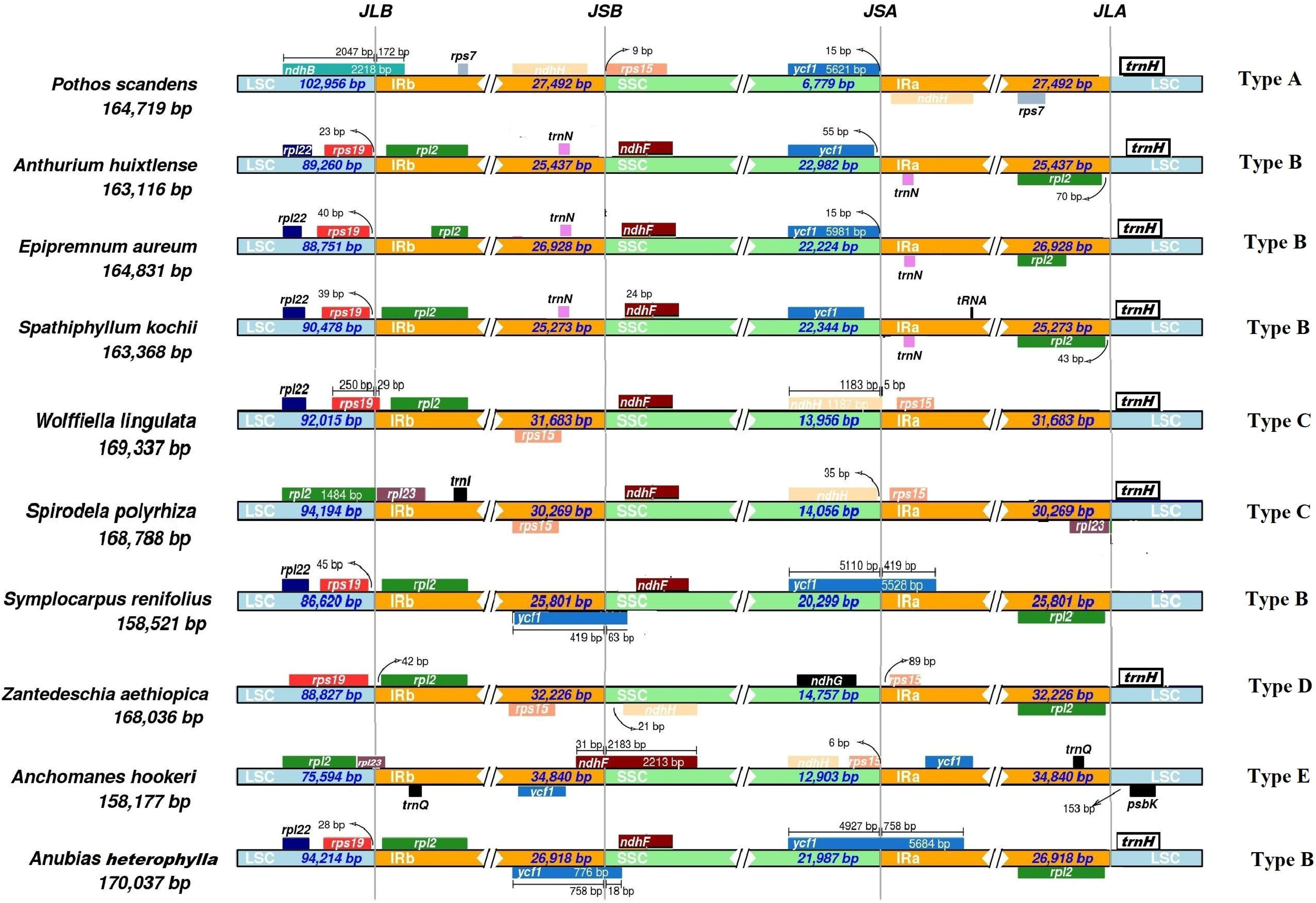
Comparative analysis of junction sites in Araceae chloroplast genomes. Abbreviations denote junction site of the plastid genome JLA (IRa/LSC), JLB (IRb/LSC), JSA (SSC/IRa) and JSB (IRb/SSC). Genes are represented by colored boxes while arrows are showing the coordinate positions of each gene near the junctions. Genes displayed on the top appear on the negative strand, while genes present bellow and found on the positive strand of the genome.

#### Effect of rate heterotachy

We also analyzed the effect of IR contraction and expansion on the evolution of protein-coding genes. Our result showed that IR contraction and expansion affects the evolution rate of protein-coding genes. The genes that were transferred from the SSC region to IR regions showed a decrease in the rate of evolution, whereas genes that travel from IR regions to the LSC region showed an increase in the rate of evolution. In *P. scandens*, we found 2454 (5.67%) substitutions in the genes of LSC, 269 substitutions (2.64%) in genes of IRs, and 1338 (9.27%) substitutions in genes of SSC. In *A. huixtlense*, we found 2428 (5.62%) substitutions in genes of LSC, 205 (2.0%) substitutions in genes of IRs, and 1327 (9.16%) in genes of SSC. We found a higher rate of evolution in *P. scandens* genes than in *A. huixtlense* and observed a difference of 0.043% in genes of LSC, 0.64% in genes of IRs, and 0.11% in genes of SSC. We observed the highest difference in evolution rate between *P. scandens* and *A. huixtlense* in IRs. This might be due to transfer of most of the IR genes of *P. scandens* to LSC region. To further verify the effect of rate heterotachy, we separately compared the rate of evolution of those genes that transferred from SSC to IRs in *P. scandens*. Genes of *P. scandens* that transferred from SSC to IRs showed 0.43% less evolution than genes of *A. huixtlense*, whereas average rate of evolution of the genes of all regions were found higher in *P. scandens* than *A. huixtlense*. This confirmed transferring of the genes from single-copy regions to IRs is responsible for decrease evolution rates.

### 3.6 Phylogenetic inference of the family Araceae

The phylogenetic tree was reconstructed with the best fit Model TVM+F+I+G4. The nucleotide alignment contained a total of 84,820 sites in which 59,644 were invariable, 11,617 were parsimony informative, and 7783 sites showed a distinct pattern. The phylogenetic tree resolved the evolutionary relationships of all species of all the subfamilies with high bootstrap support (Fig. 6), apart from the position of *Epipremnum aureum*. The *Epipremnum* genus is included in subfamily Monsteroideae (Zuluaga et al. 2019) but here was found to be embedded in subfamily Aroideae and to share a node with *Zantedeschia*. This result was previously observed by Kim et al. (2019) and Henriquez et al. (2020a). This might be due to chloroplast capture by this species of some other species or due to misidentification of the species.

**Fig 6.**
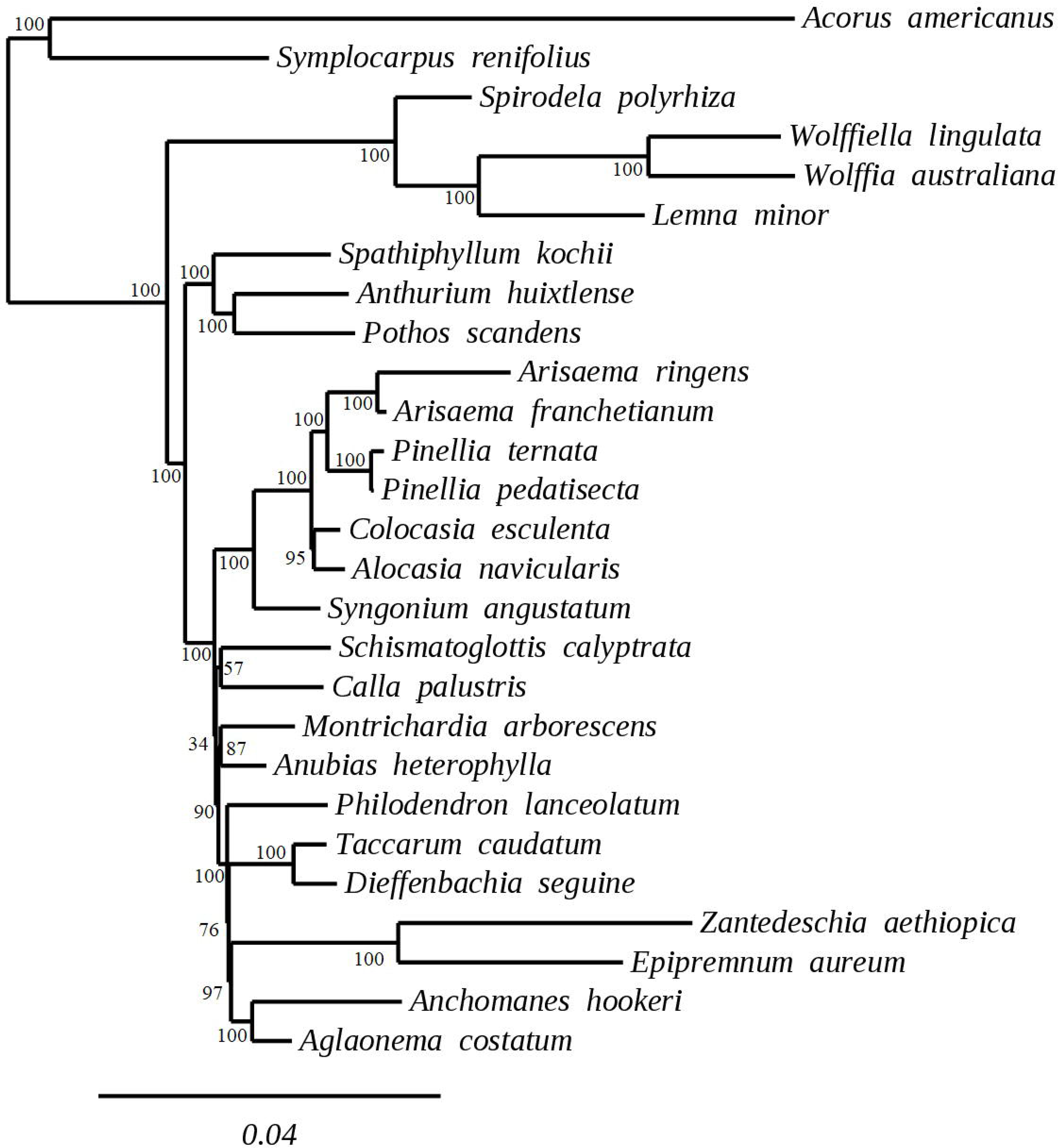
Maximum likelihood phylogenetic tree of the Araceae family reconstructed from plastid genome data

## Discussion

In the current study, we assembled the chloroplast genomes of two species from the subfamily Pothoideae. The chloroplast genomes of both *P. scandens* and *A. huixtlense* were found to be unique among Araceae species and showed a unique type of IR contraction and expansion that affected the evolution rate in *P. scandens*.

In the current study, the chloroplast genome of *P. scandens* showed uneven IR contraction and expansion, which led to a variable number of genes. IR contraction and expansion is very common in chloroplast genomes and leads to variation in the number of genes in various plant lineages, including Araceae (Menezes et al. 2018; Cho et al. 2018; Lee et al. 2018; Abdullah et al. 2020; Henriquez et al. 2020a). IR contraction and expansion also results in new combinations of genes in the IR regions, which in turn leads to rearrangement of genes in the SSC region, as previously reported in Araceae (Wang and Messing 2011; Henriquez et al. 2020a). However, in *P. scandens* we observed the formation of a new combination of genes in IRs but not an accompanying rearrangement of the genes. A similar effect of IR contraction and expansion was also reported in other plant lineages without any effect on the arrangement of genes (Cho et al. 2018; Lee et al. 2018). In *P. scandens*, the SSC region showed significant shortening and contained only two genes. Similar shortening of the SSC region was also reported in other angiosperms and even smaller SSC regions have been reported (Cho et al. 2018; Lee et al. 2018). Previously, four types of gene arrangements were observed in Araceae. Two types of gene arrangements were observed at IR junctions in one comparison of Araceae species (Choi et al. 2017) and two other types of gene arrangements at the junctions were reported in the chloroplast genomes of two species of subfamily Aroideae, including *Anchomanes hookeri* and *Zantedeschia aethiopica* (Henriquez et al. 2020a). In the current study, we report a fifth type of gene arrangement at the junctions in the chloroplast genome of *P. scandens*. Further genomic resources from the genus *Pothos* and subfamily Pothoideae might be helpful to gain insight into chloroplast genome structure and to discern whether this uneven IR contraction and expansion occurs only in *P. scandens* or in the genus *Pothos* as a whole.

The expansion of IR regions in our study decrease the evolution rate of protein-coding genes that transfer from SSC to IR, whereas an increase in the evolution rate can be observed in the genes that transfer from LSC to IRs. Similar results were reported in the chloroplast genomes of other species and a higher rate was observed in the regions that exist in the single-copy region instead of IR region (Zhu et al. 2016). In contrast, the effect on evolution rate in *Pelargonium* was not observed due to IR contraction and expansion (Weng et al. 2017). The low evolution rate of genes that exist in IR regions might be due to a repairing mechanism (Zhu et al. 2016).

We observed high GC content in the IR regions when compared with the LSC and SSC regions. Chloroplast genomes are mostly conserved in terms of gene content and organization, and similar observations were also reported in other angiosperms including other subfamilies of Araceae (Wang and Messing 2011; Iram et al. 2019; Abdullah et al. 2020; Henriquez et al. 2020b, a; Shahzadi et al. 2020). The IR regions of the genome of *P. scandens* showed a decrease in GC content up to 5% when compared with *A. huixtlense*. This might be due to expansion of the IR regions, which leads to inclusion of most of the SSC region (which has low GC content).

Amino acid frequency analyses showed a high encoding frequency of leucine and iso-leucine and a low frequency of cysteine. Higher RSCU values (≥ 1) were found for codons with A or T at the 3’ position and showed high encoding efficacy. Similar results for amino acid frequency and codon usage have also been reported in the chloroplast genomes of other angiosperms, which might be due to the high AT content in the chloroplast genome (Amiryousefi et al. 2018b; Menezes et al. 2018; Abdullah et al. 2019b; Mehmood et al. 2020). The analyses of oligonucleotide repeats showed the existence of four types of repeats, but the repeats varied in size and types between the two species. The variation in the types and size of repeats were also previously reported in the chloroplast genomes of angiosperms and in other species of Araceae (Abdullah et al. 2020; Mehmood et al. 2020; Henriquez et al. 2020a). These repeats can be used as a proxy to identify mutational hotspots (Ahmed et al. 2012).

We found higher levels of transition substitutions than transversion substitutions. This is expected in all types of DNA sequences (Wakeley 1996). However, fewer transition substitution have been reported when compared to transversion substitutions in chloroplast genomes (Cai et al. 2015; Abdullah et al. 2019a; Shahzadi et al. 2020). This bias of higher transversions might be due to the composition of genomes and the genetic characteristics of codons (Morton et al. 1997). Our study is consistent with the statement above regarding the characteristics of codons, as we observed higher transition substitutions in synonymous substitutions than non-synonymous substitutions. A similar result was also reported in the chloroplast genomes of *Firmiana*, a genus of family Malvaceae (Abdullah et al. 2019a). Here, we reported a higher rate of transitions in the coding genes than that of transversions. Similar results were also reported in the coding sequences of the species of Lemnoideae (Araceae) and in the complete chloroplast genome of *Dioscorea polystachya* (Cao et al. 2018).

The rate of evolution of protein coding genes showed strong purifying selection pressure. The higher rate of synonymous substitutions than non-synonymous substitutions suggests strong purifying selection pressure acting on these genes during the course of evolution (Matsuoka et al. 2002). These results are also consistent with previous studies on angiosperm chloroplast genomes and with the closest subfamily (Monsteroideae) of Pothoideae (Menezes et al. 2018; Abdullah et al. 2019a, 2020; Henriquez et al. 2020b; Shahzadi et al. 2020). Here, no protein-coding genes were found under positive selection pressure. However, seven genes in Monsteroideae were found under positive selection pressure, whereas in another study of Araceae most genes were reported under positive selection pressure (Kim et al. 2019).

In conclusion, our study provides insight into the evolution of chloroplast genomes of Pothoideae (Araceae). Our study shows unique IR contraction and expansion affecting the number of genes and rate of evolution in *P. scandens*. We observed a two-fold higher transition substitution rate than transversion substitutions and found higher transversion substitutions linked with non-synonymous substitutions when compared with transition substitutions.

## Acknowledgment

The authors would like to thank Dr. Barbara Schaal at Washington University in St. Louis and Dr. J. Chris Pires at the University of Columbia, Missouri for funding and laboratory support. The authors would also like to thank Dr. Tatiana Arias for valuable help in the laboratory and data processing. The authors also thank Emily Colletti in the aroid greenhouse at the Missouri Botanical Garden for help with living material.

